# Legal but immoral: attitudes toward non-invasive brain stimulation for cognitive enhancement

**DOI:** 10.1101/2025.09.08.674869

**Authors:** M. Hurford, A. Dunham, S. Selvaratnam, A. K. Martin

## Abstract

Cognitive enhancement involves using substances or technologies to improve mental performance in healthy individuals. While methods such as caffeine are widely accepted, others, including prescription stimulants and non-invasive brain stimulation (tDCS), evoke ethical concern. This study examined how university students evaluate three forms of cognitive enhancement (natural, pharmacological, brain stimulation) across five moral domains: academic fairness, free will, naturalness, self-identity, and safety. We also tested how evaluations were shaped by framing (preservation vs. enhancement) and priming (self-affecting vs. non-self-affecting). A total of 449 students completed an online experiment with a 2 × 2 × 3 mixed design, rating enhancement acceptability after reading intervention-specific scenarios. Natural enhancers were judged most positively, followed by brain stimulation and then pharmacological agents. Importantly, although brain stimulation is legal and marketed as non-invasive, it was evaluated more like pharmacological enhancement than natural methods, indicating that legality does not equate to moral acceptability. Framing effects showed that participants were more accepting of enhancement when it was described as preserving ability rather than augmenting it beyond baseline. Priming participants to consider how others’ use might affect their own outcomes reduced moral acceptability, particularly for pharmacological and brain stimulation methods in the domain of academic fairness. Overall, public attitudes toward enhancement appeared shaped less by legality or objective safety than by intuitive moral reasoning. These findings highlight the importance of adopting pluralistic, context-sensitive approaches to policy and regulation, as student judgments reflect concerns about fairness, authenticity, and competition in academic life.

## Introduction

Cognitive enhancement is the use of techniques or substances to improve cognitive performance in healthy individuals and has become a key focus in debates across neuroscience, bioethics, and education. With growing availability of both pharmacological and non-pharmacological tools aimed at improving cognitive abilities such as memory, attention, and executive functioning, individuals are increasingly presented with the option to modify cognition, not just in a clinical context, but in everyday life. These tools range from commonplace and widely accepted substances such as caffeine, to prescription-only pharmaceuticals like modafinil and methylphenidate (Ritalin), to newer, non-invasive brain stimulation technologies such as transcranial direct current stimulation (tDCS). While these methods vary in mechanism, accessibility, and risk, they all raise normative questions about what kinds of enhancement are acceptable, and under what conditions.

University students form a particularly relevant population for studying cognitive enhancement. As a group under consistent academic pressure and performance scrutiny, students may be both consumers of enhancement technologies and critical evaluators of their legitimacy. Existing data suggest increasing interest in the use of prescription stimulants such as modafinil among UK students without medical need (e.g., McDermott et al., 2021). Yet despite rising use, student attitudes toward enhancement are complex and context-sensitive. While some enhancers are viewed as harmless or even necessary for coping with academic life (Franke et al., 2011), others are seen as morally questionable, potentially unfair, or indicative of an unhealthy culture of overperformance (Maier et al., 2015).

A growing body of literature has examined the factors that shape public judgments of cognitive enhancement. A key finding is that attitudes are not driven solely by considerations of efficacy or legality, but are influenced by underlying moral intuitions, particularly those related to fairness and authenticity (Caviola & Faber, 2015; Fitz et al., 2014; Scheske & Schnall, 2012). For example, people are generally more accepting of enhancement when it is used to restore or preserve abilities, and more sceptical when it is used to push cognitive performance beyond the perceived baseline (Maslen et al., 2014; Medaglia et al., 2019). This distinction between treatment and enhancement plays a powerful role in shaping moral reasoning, although the boundary between treatment and enhancement is far from distinct (Maslen et al., 2014). It becomes even more opaque when moralising over temporary states of impairments often faced by students and workers, such as stress or tiredness.

In addition to concerns about fairness and authenticity, cognitive enhancement also raises questions of identity. Scholars have argued that altering cognitive capacities can be seen as altering the self, threatening continuity of personal identity or undermining the authenticity of one’s achievements (Parens, 2005; Schermer, 2008). Relatedly, safety concerns often shape public debates, not only in terms of objective health risks but also in terms of perceived legitimacy: interventions judged unsafe are more easily construed as morally problematic (Cabrera & Reiner, 2015). Although caffeine is widely used despite recognised risks (Costantino et al., 2023), newer technologies such as brain stimulation provoke unease precisely because their safety is relatively unknown amongst the general population (Furrer et al., 2025). Together, these themes suggest that judgments about enhancement reflect not just efficacy or legality, but also deeper intuitions about what it means to act authentically, preserve identity, and protect oneself and others from harm.

Social context also matters. Dinh et al. (2020) demonstrate that attitudes toward pharmacological enhancement shift depending on who is using it and under what conditions. People tend to be more permissive when enhancement is used by themselves or by peers in low-stakes situations, but become more critical when it is used by others in competitive contexts. In such cases, the moral concern is less about the intrinsic nature of the enhancer and whether it is natural, legal, or medically safe and more about its perceived impact on fairness and merit. Specifically, enhancement is seen as ethically problematic when it is believed to give others an unfair advantage, disrupt equal opportunity, or undermine the legitimacy of performance outcomes. This suggests that moral evaluations are not only about the enhancer itself, but also about who benefits from it, and in what social context. This aligns with research suggesting that cognitive enhancement is evaluated through a relational lens, where judgments are sensitive to power dynamics, competitive settings, and concerns over social disruption (Conrad et al., 2019; Forlini & Racine, 2009).

The type of enhancement method also plays a role in shaping attitudes. While natural substances like caffeine are broadly accepted (Franke et al., 2011), pharmacological agents such as modafinil and methylphenidate are often met with ethical concern, particularly when used *illegally* without prescription (Schelle et al., 2014). Meanwhile, brain stimulation techniques like tDCS are legal and marketed as non-invasive, yet remain unfamiliar to many and may evoke ambivalence or unease. However, research comparing ethical judgements on cognitive enhancement via brain stimulation, in comparison to pharmacological or natural methods, has remained unexplored. Veit et al (2020) argue the importance of recognising the diversity of cognitive enhancement methods in relation to ethical concerns. Rather than adopting one-size-fits-all ethical frameworks, researchers and policymakers must consider how different tools interact with different values, contexts, and user intentions. Ethical analysis must move beyond blanket judgments and instead adopt a pluralistic, context-aware perspective.

The present study responds to this call by exploring how university students evaluate three forms of cognitive enhancement—natural substances (e.g., caffeine), pharmacological agents (e.g., modafinil, methylphenidate), and brain stimulation (e.g., tDCS)—across five moral domains: academic fairness, free will, self-identity, naturalness, and safety. In addition to comparing these methods, we test how attitudes shift depending on framing (preservation vs. enhancement) and priming (self-relevant vs. non-self-relevant scenarios). By incorporating these contextual dimensions, we aim to provide a more fine-grained understanding of how moral intuitions shape judgments about enhancement in real-world academic environments.

## Method

### Participants

Participants were recruited through convenience sampling using the University of Kent School of Psychology Research Participation Scheme (RPS). Eligible participants were required to be enrolled university students with fluent English comprehension. To ensure data quality, individuals who failed one or more attention checks or failed to complete the study were removed. This resulted in the removal of 81 who failed to complete the entire questionnaire. None of the participants failed the attention check. The final sample comprised 449 participants, which exceeded the minimum sample size (N=240) required to detect a small effect (f = 0.10) with 90% power at α =.05, as determined using G*Power (Faul et al., 2007).

### Design and Measures

This study employed a 2 (priming condition: self-affecting vs. non-self-affecting) × 2 (framing: preservation vs. enhancement) × 3 (intervention type: pharmacological, brain stimulation, natural) mixed-factorial design. Priming condition and framing were between-subject variables, while intervention type was a within-subject variable. The primary dependent variable was attitude toward cognitive enhancement, operationalised through theme-specific mean scores across five moral domains: fairness in academia, autonomy and free will, self-identity, safety, and naturalness.

Attitudes were measured using 20 items adapted from the Attitudes Toward Pharmacological Cognitive Enhancement Scale (Patton, 2023), which has previously demonstrated high internal consistency (α =.80). Each item was rated on a 7-point Likert scale ranging from -3 (“Strongly disagree”) to +3 (“Strongly agree”), with 0 indicating neutrality. Items were reverse-coded where necessary to ensure consistency in directionality, with higher scores indicating greater acceptability of enhancement.

### Procedure

Participants completed the study online via Qualtrics. After reading an information sheet and providing informed consent, participants generated an anonymous code to protect identity and enable withdrawal if requested. They first completed demographic questions and a brief set of items measuring prior awareness and use of cognitive enhancement techniques (see Supplementary 1).

Participants were then randomly assigned to one of two priming conditions (see Supplementary 1). In the self-affecting condition, the scenario stated that other students’ grades could impact the participant’s own academic outcomes. In the non-self-affecting condition, the scenario suggested that others’ performance had no impact on the participant’s own results. Both primes were followed by a 15-second countdown and two attention check items.

Following priming, participants were randomly assigned to one of two framing conditions (see Supplementary 1), which described each intervention (natural enhancers, brain stimulation, and pharmaceuticals) as either “preserving” cognitive ability or “enhancing” it. Participants read three scenarios (one per intervention type) presented in randomised order. Each scenario was followed by 20 questions (four per theme), completed on the same Likert scale. The full survey took approximately 25 minutes to complete. Upon completion, participants were debriefed and awarded course credit as part of their undergraduate degree programme.

### Ethical Considerations

This study was reviewed and approved by the University of Kent Research Ethics Committee [Ref: 202417314953719401]. All procedures conformed to the British Psychological Society’s Code of Ethics and Conduct.

Participants were fully informed of their rights and could withdraw at any point. No deception was involved, and although the study posed no physical risks, some items referred to potentially sensitive topics such as the use of controlled substances or perceived academic pressure. To minimise discomfort, responses were anonymous and non-identifiable, and contact details for the research team and ethics committee were provided in case of concerns.

All data were stored securely on password-protected university servers, with access restricted to the research team. In accordance with the General Data Protection Regulation (GDPR), only minimal personally identifiable information was collected.

### Statistical Analysis

Data were analysed using JASP (Version 0.19.3; JASP Team, 2025). For each of the five moral themes, a 2 (Priming) × 2 (Framing) × 3 (Enhancement Type) mixed repeated measures ANOVA was conducted to assess main effects and interactions. Planned contrast analyses were conducted to test specific hypotheses regarding perceptions of enhancement type. The contrast weights used were: natural = 1, brain stimulation = -2, and pharmaceutical = 1, allowing for direct comparison between natural and pharmaceutical enhancers relative to brain stimulation. When appropriate, post hoc pairwise comparisons with Holm correction were used to examine differences between enhancement types.

## Results

### Demographics

The final sample consisted of 449 participants. After exclusions, the groups were slightly imbalanced: Self-affecting/Preservation (N = 106), Self-affecting/Enhancement (N = 107), Non-self-affecting/Preservation (N = 119), and Non-self-affecting/Enhancement (N = 117). The average age was 19.90 years (SD = 3.85; range = 18–54 years). The majority identified as female (81.7%), with 15.6% identifying as male, 2.2% as third gender or non-binary, and 0.4% preferring not to disclose. In terms of ethnicity, 64.0% identified as White, 15.1% as Black/African/Caribbean/Black British, 12.1% as Asian/Asian British, 4.0% as Mixed or multiple ethnic groups, and 3.3% as Other, while 1.1% preferred not to say.

92% of participants were aware of natural cognitive enhancers, 83.3% were aware of pharmacological cognitive enhancers, and 39.0% aware of brain stimulation as a cognitive enhancement tool. Ten participants decided not to disclose their use of cognitive enhancers. Of the remainder (N=439), 298 (67.88%) reported using natural substances, 28 (6.38%) reported using brain stimulation, and 40 (9.11%) reported using illegal pharmacological methods, to enhance cognitive functioning.

All mean attitude scores for each cognitive enhancement method across each moral theme are presented in Table 1.

**Table 1.** Attitudes towards natural, brain stimulation, and pharmacological enhancers across the moral themes of academic fairness, free-will, naturalness, self-identity, and safety

|  | Natural | Brain Stimulation | Pharmacological |
| --- | --- | --- | --- |
|  | Mean (sd) | Mean (sd) | Mean (sd) |
| Academia |  |  |  |
| *Self-affecting* |  |  |  |
| Preservation | 1.54 (1.12) | 0.11 (1.08) | -0.13 (1.16) |
| Enhancement | 1.44 (1.10) | -0.11 (1.25) | -0.47 (1.18) |
| *Non self-affecting* |  |  |  |
| Preservation | 1.57 (1.04) | 0.28 (1.13) | 0.12 (1.22) |
| Enhancement | 1.44 (1.02) | 0.22 (1.10) | -0.06 (1.15) |
| Free-will |  |  |  |
| *Self-affecting* |  |  |  |
| Preservation | 1.36 (0.96) | 0.62 (0.95) | 0.44 (1.12) |
| Enhancement | 0.95 (0.94) | 0.41 (1.00) | 0.19 (1.01) |
| *Non self-affecting* |  |  |  |
| Preservation | 1.02 (1.12) | 0.52 (1.15) | 0.33 (1.18) |
| Enhancement | 1.13 (1.09) | 0.67 (1.11) | 0.39 (1.16) |
| Naturalness |  |  |  |
| *Self-affecting* |  |  |  |
| Preservation | 0.93 (0.93) | 0.14 (0.90) | 0.01 (0.90) |
| Enhancement | 0.81 (0.96) | -0.10 (0.91) | -0.22 (0.86) |
| *Non self-affecting* |  |  |  |
| Preservation | 0.84 (0.88) | 0.06 (0.92) | -0.05 (0.96) |
| Enhancement | 0.87 (0.99) | 0.10 (0.98) | -0.11 (0.97) |
| Self-Identity |  |  |  |
| *Self-affecting* |  |  |  |
| Preservation | 1.51 (1.03) | 0.74 (1.00) | 0.38 (1.09) |
| Enhancement | 1.40 (1.09) | 0.53 (1.02) | 0.23 (1.02) |
| *Non self-affecting* |  |  |  |
| Preservation | 1.47 (1.00) | 0.70 (1.13) | 0.44 (1.13) |
| Enhancement | 1.47 (1.02) | 0.73 (1.09) | 0.40 (1.07) |
| Safety |  |  |  |
| *Self-affecting* |  |  |  |
| Preservation | 0.59 (0.72) | 0.57 (0.62) | 0.51 (0.58) |
| Enhancement | 0.57 (0.67) | 0.42 (0.51) | 0.47 (0.52) |
| *Non self-affecting* |  |  |  |
| Preservation | 0.49 (0.70) | 0.44 (0.62) | 0.49 (0.53) |
| Enhancement | 0.52 (0.74) | 0.56 (0.56) | 0.42 (0.59) |

*Note.* All themes were scored from -3 (negative) to +3 (positive)

### Academic Fairness

The main effects of Enhancement Type, *F*(1.74, 773.71) = 463.61, *p* <.001, *η*^*2*^_*p*_ = 0.51, Framing, *F*(1, 445) = 4.29, *p* =.04 (see Figure 1), *η*^*2*^_*p*_ = 0.01, and Priming, *F*(1, 445) = 5.71, *p* =.02, *η*^*2*^_*p*_ = 0.01, were significant. A post-hoc pairwise comparison using Holm correction showed that Natural methods were rated significantly higher than Brain Stimulation, t(445) = 23.81, *p* <.001, d=1.21 and Pharmacological agents, t(445) = 28.34, *p* <.001, d=1.45. Brain Stimulation was rated significantly higher than Pharmacological agents, t(445) = 4.53, *p* <.001, d=0.23. The contrast between Natural and Brain Stimulation and Brain Stimulation and Pharmacological agents was significant, *t*(890) = 11.13, *p* <.001 (see Figure 2).

**Figure 1.**
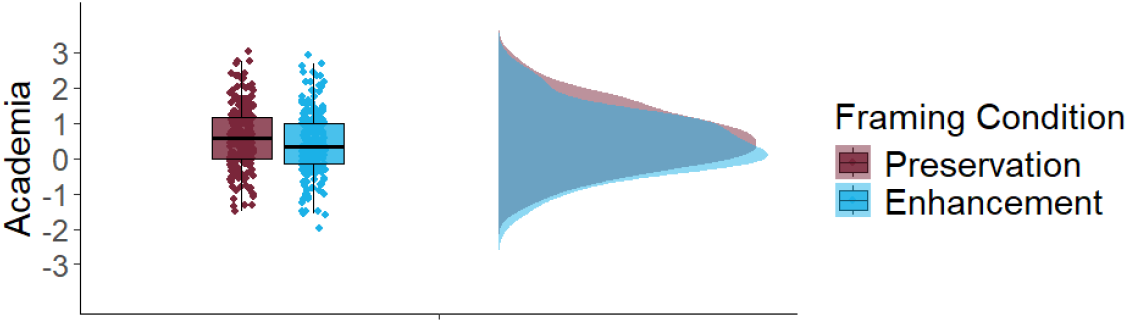
Academic fairness attitudes towards cognitive enhancers were more positive in the preservation scenario across all enhancement types and priming conditions

**Figure 2.**
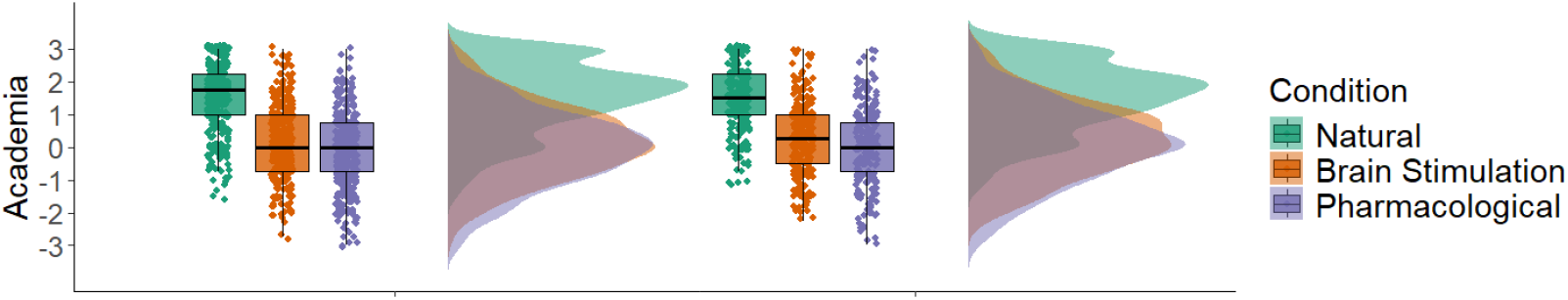
Academic fairness attitudes showed a similar pattern across self-affecting priming conditions for natural enhancers but attitudes were not as positive for brain stimulation and pharmacological enhancers when others enhancing could affect their grades

A priming x enhancement type interaction was also identified, F(1.74, 773.71)=3.90, p=.03, *η*^*2*^_*p*_ =0.01. Post-hoc t-tests showed a significant effect of priming for both the pharmacological enhancement, t(447)=2.97, p=.003, d=0.28, and brain stimulation, t(447)=2.31, p=.02, d=0.22, but not natural, t(447)=0.21, p=.84, d=0.02.

### Free will

The main effect of Enhancement Type, *F*(1.82, 811.23) = 183.83, *p* <.001, *η*^*2*^_*p*_ = 0.29 was significant.

Natural methods were rated significantly higher than Brain Stimulation, *t*(445) = 13.40, *p* <.001, *d* = 0.52) and Pharmacological agents, *t*(445) = 18.58, *p* <.001, *d* = 0.72). Brain Stimulation was also rated significantly higher than Pharmacological agents, *t*(445) = 5.18, *p* <.001, *d* = 0.20; see Figure 3.

**Figure 3.**
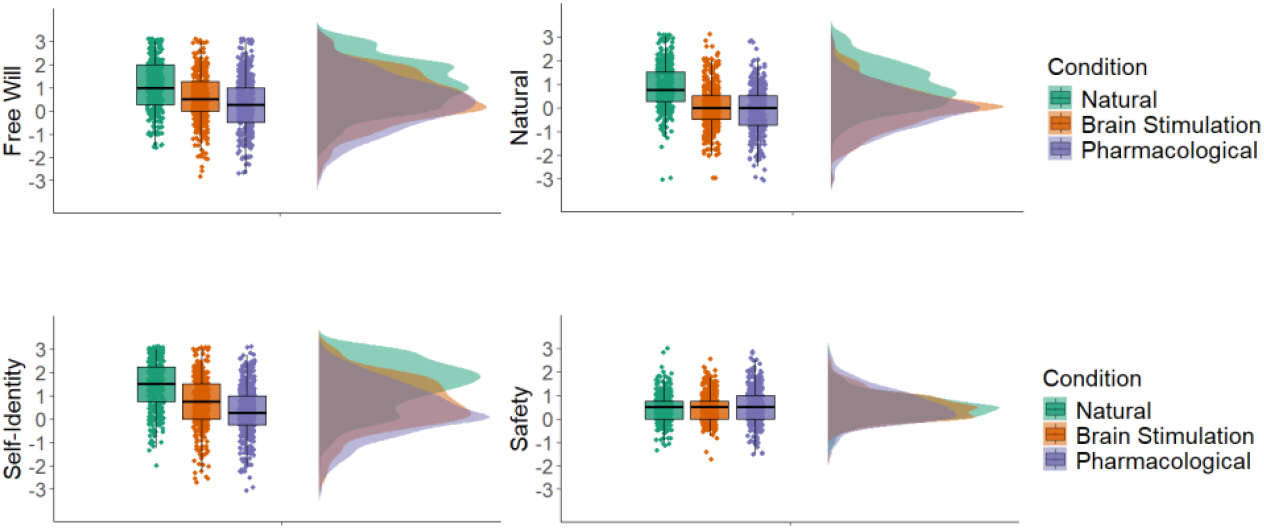
Student attitudes towards free will, naturalness, self-identity, and the safety of pharmacological, brain stimulation, or natural enhancers. *Note*. All enhancement conditions differed significantly from one another, except for Safety, where no significant differences were observed between natural, brain stimulation, and pharmacological enhancers

The contrast between Natural and Brain Stimulation and Brain Stimulation and Pharmacological agents was significant, *t*(890) = 4.75, *p* <.001.

The main effects of Framing, *F*(1, 445) = 1.09, *p* =.30, η^2^p =.002, and Priming, *F*(1, 445) = 0.03, *p* =.86, η^2^p <.001, were both non-significant. However, the Priming × Framing interaction was significant, *F*(1, 445) = 4.95, *p* =.03, η^2^p =.01. Follow-up Holm-corrected comparisons did not identify any reliable simple effects. Within each priming condition (self-affecting vs. non-self-affecting), responses did not differ significantly between preservation and enhancement framing. Likewise, within each framing condition, the two priming groups did not differ significantly from one another (all *p*s >.14).

The interaction effects between Enhancement Type and Framing condition, *F*(1.82, 811.23) = 1.06, *p* =.34, *η*^*2*^_*p*_ = 0.002 or Priming condition, *F*(1.82, 811.23) = 2.17, *p* =.12, *η*^*2*^_*p*_ = 0.005 were non-significant. Additionally, the three-way interaction was non-significant, *F*(1.82, 811.23) = 0.96, *p* =.38, *η*^*2*^_*p*_ =0.002.

### Naturalness

The main effect of Enhancement Type, *F*(1.76, 784.53) = 258.88, *p* <.001, *η*^*2*^_*p*_ = 0.37, was significant, suggesting that attitudes differed significantly across methods. However, the main effects of Framing, *F*(1, 445) = 1.80, *p* =.18, *η*^*2*^_*p*_ = 0.004, and Priming, *F*(1, 445) = 0.13, *p* =.72, *η*^*2*^_*p*_ < 0.001, were non-significant.

A post-hoc pairwise comparison using Bonferroni correction showed that Natural methods were rated significantly higher than brain stimulation, t(445) = 17.91, *p* <.001, d=0.87, and pharmacological agents, t(445) = 21.11, *p* <.001, d=1.02). Brain Stimulation was rated significantly higher than Pharmacological agents, t(445) = 3.21, *p* =.001, d=0.16 (see Figure 3).

The contrast between Natural and Brain Stimulation and Brain Stimulation and Pharmacological was significant, *t*(445) = 8.46, *p* <.001.

The interaction effects between Enhancement Type and Framing condition, *F*(1.76, 784.53) = 0.62, *p* =.52, *η*^*2*^_*p*_ = 0.001 or Priming condition, *F*(1.76, 784.53) = 0.31, *p* =.71, *η*^*2*^_*p*_ < were non-significant. Additionally, the three-way interaction was non-significant, *F*(1.76, 784.53) = 0.33, *p* =.69, *η*^*2*^_*p*_ = 0.001, suggesting that the relationship between Priming, Framing and Enhancement Types did not significantly interact to influence attitudes of naturalness.

### Self-Identity

There was a significant main effect of Enhancement Type, *F*(1.81, 807.02) = 256.94, *p* <.001, η^2^p =.366, indicating that attitudes differed depending on the type of cognitive enhancement. Pairwise comparisons showed that natural methods were rated more positively than brain stimulation, t(445) = 15.77, *p* <.001, d=0.74 and pharmacological agents, t(445) = 21.99, *p* <.001, d=1.04. Brain stimulation was rated more positively than pharmacological agents, t(445)= 6.22, *p* <.001, d=0.29 (see Figure 3).

By contrast, there were no significant main effects of Priming, *F*(1, 445) = 0.70, *p* =.40, η^2^p =.002, or Framing, *F*(1, 445) = 0.92, *p* =.338, η^2^p =.002. Likewise, none of the interactions were significant: Enhancement × Priming, *F*(1.81, 807.02) = 0.51, *p* =.584, η^2^p =.001; Enhancement × Framing, *F*(1.81, 807.02) = 0.11, *p* =.882, η^2^p <.001; Priming × Framing, *F*(1, 445) = 0.90, *p* =.343, η^2^p =.002; or Enhancement × Priming × Framing, *F*(1.81, 807.02) = 0.31, *p* =.715, η^2^p <.001.

Post hoc comparisons with Holm correction showed that natural methods were rated more positively than brain stimulation, *t*(445) = 15.77, *p* <.001, and pharmacological agents, *t*(445) = 21.99, *p* <.001. Brain stimulation was also rated more positively than pharmacological agents, *t*(445) = 6.22, *p* <.001.

The contrast between Natural and Brain Stimulation and Brain Stimulation and Pharmacological was significant, *t*(890) = 5.51, *p* <.001, indicating greater overlap between brain stimulation and pharmacological enhancers.

The interaction effects between Enhancement type and Framing condition, *F*(1.83, 972.80) = 0.50, *p* =.590, *η*^*2*^_*p*_ < 0.01, or Priming condition, *F*(1.83, 972.80) = 0.36, *p* =.678, *η*^*2*^_*p*_ < 0.01 were non-significant. Additionally, the three-way interaction was non-significant, *F*(1.83, 972.80) = 0.67, *p* =.498, *η*^*2*^_*p*_ < 0.01, suggesting the relationship between Priming, Framing and Enhancement Types did not significantly interact to influence attitudes pertaining to the self.

### Safety

The main effects of Enhancement Type, *F*(1.91, 850.20) = 2.28, *p* =.11, *η*^*2*^_*p*_ = 0.01, Framing, *F*(1, 445) = 0.20, *p* =.66, *η*^*2*^_*p*_ < 0.001, and Priming, *F*(1, 445) = 0.75, *p* =.39, *η*^*2*^_*p*_ < 0.002 were non-significant.

The interaction effects between Enhancement Type and Framing condition, *F*(1.91, 850.20) = 0.44, p=.54, *η*^*2*^_*p*_ = 0.001, or Priming condition, *F*(1.91, 850.20) = 0.78, *p* =.45, *η*^*2*^_*p*_ = 0.002 were non-significant. Additionally, the three-way interaction was non-significant, *F*(1.91, 850.20) = 3.00, *p* =.053, *η*^*2*^_*p*_ = 0.007, suggesting the relationship between Priming, Framing and Enhancement Types did not significantly interact to influence student attitudes concerning safety.

## Discussion

We assessed student attitudes to cognitive enhancers under varying contextual conditions. Our study contributes to a growing body of research exploring the social and moral psychology of cognitive enhancement (Grinschgl et al., 2025), offering new insights into how students evaluate different enhancement methods under varying contextual conditions. One of our central findings is that brain stimulation, although legal and non-invasive, was judged more similarly to pharmacological enhancement than to natural methods such as caffeine. This indicates that legal status alone does not guarantee moral acceptance, similar to previous findings (Mayor et al., 2019; Mihailov et al., 2021). Instead, participants appeared to rely on deeper, intuitive judgments about the naturalness or familiarity of different enhancement types. This is consistent with prior work suggesting that people evaluate brain stimulation with caution due to its perceived artificiality or invasiveness (Cohen Kadosh et al., 2012), even in the absence of clinical risk (Bikson et al., 2016).

Our results align closely with Dinh et al. (2020), who found that moral concern about cognitive enhancement often arises not from objective characteristics of the enhancer itself, but from its implications for fairness and social comparison. In our study, students perceived brain stimulation as occupying a middle ground between natural and pharmacological enhancers, but importantly, with closer alignment to pharmacological enhancers, despite its legal and commercially available status. This suggests that students are forming categories not based on legality or availability, but on perceived moral distance from socially accepted practices.

Self-identity concerns also emerged as a central dimension of students’ moral judgments. Whereas natural enhancers were largely seen as compatible with a stable sense of self, both pharmacological and brain stimulation methods were more likely to evoke unease about authenticity and continuity of identity. These findings align with the idea that enhancement is not evaluated solely in terms of individual choice but in relation to implicit concerns about social equity and moral status. Francis Fukuyama’s (2002) notion of “Factor X,” a core human essence grounding dignity, offers a useful lens: when cognitive capacities are seen as modifiable through artificial means, this may be perceived as undermining the shared basis of moral equality. In competitive academic contexts, threats to selfhood are not only experienced individually but also carry relational implications: if some students are perceived to enhance in ways that compromise authenticity, this can destabilise the sense of equal moral standing that underpins fair competition. In this way, identity-related discomfort intersects with fairness concerns, reinforcing unease about legitimacy and equity in academic achievement.

These findings reinforce Veit et al’s (2020) call to recognise the diversity of cognitive enhancement in ethical discussions. Rather than treating enhancement as a monolithic category, our data show that students distinguish between methods in nuanced ways, with attitudes shaped by perceived invasiveness, familiarity, and social meaning. Brain stimulation, despite being new and legally available, does not benefit from the same moral leniency as natural enhancers, and instead inherits the ethical burden often placed on pharmacological agents. A pluralistic, context-sensitive ethical framework, one that evaluates each form of enhancement on its own terms, is therefore essential for capturing how these technologies are understood and morally negotiated.

Another key finding relates to the effect of framing. Across all enhancement types, participants expressed more favourable attitudes when the enhancer was described as preserving or restoring cognitive function, rather than improving it beyond baseline. This finding supports the well-established treatment–enhancement distinction in moral psychology (Maslen et al., 2014; Medaglia et al., 2019). Notably, our scenarios involved only minor and non-clinical deficits, such as tiredness or lack of focus, yet the preservation framing still shifted responses. This indicates that the treatment framing may serve as a moral justification mechanism, making enhancement feel less like an unfair advantage and more like a form of responsible self-care. These results also align with Dinh et al. (2020), who found that people tend to accept enhancement more readily when it is framed as addressing a need rather than augmenting ability.

Our priming manipulation, which varied whether enhancement was framed as self-relevant, also significantly shaped moral judgments. When participants were asked to consider how others’ use of enhancement could affect their own academic outcomes, they evaluated enhancement more negatively, particularly for pharmacological and brain stimulation methods. In contrast, judgments of natural enhancers remained relatively stable. The effect was especially pronounced in themes related to fairness and identity, suggesting that enhancement is not merely a personal choice but a socially consequential one. These findings are consistent with research showing that enhancement in zero-sum environments (for example, competitive academic contexts) is judged more harshly, as it may threaten perceptions of merit and equity (Fitz et al., 2014; Scheske & Schnall, 2012). Our results extend this by showing that even mild priming, which highlights how others’ actions might impact one’s own standing, can activate fairness-related moral concerns, but primarily in relation to pharmacological and neural interventions. One possible explanation is that these methods are seen as more artificial and more capable of producing substantial performance gains, which heightens concerns about undeserved advantage, authenticity, and social comparison. Natural enhancers, by contrast, may be perceived as part of everyday academic life and therefore less threatening to fairness or identity.

Interestingly, safety judgments did not differ significantly between natural, pharmacological, and brain stimulation methods, in contrast to previous findings (Cabrera & Reiner, 2015; Schelle et al., 2014). One explanation is that participants may have had limited or inconsistent knowledge about the objective risks of different enhancers, leading to broadly similar evaluations across categories. For example, although caffeine is widely used, it can carry recognised health risks such as dependence, anxiety, and sleep disruption (Franke et al., 2011). By contrast, the safety of prescription stimulants such as methylphenidate and modafinil remains debated, particularly when used without medical supervision (Schelle et al., 2014). Brain stimulation techniques such as tDCS are considered safe under controlled research and clinical conditions, but questions remain about unsupervised or commercial use (Bikson et al., 2016). This effect may also reflect the characteristics of our sample, which consisted primarily of psychology students. Such students are regularly exposed to training and discussions around research methods and participant protection, which may foster a generalised sensitivity to safety concerns. As a result, they may have adopted a cautious but relatively undifferentiated stance across enhancer types rather than privileging one method as inherently safer than another. In contrast, domains such as fairness or identity may rely less on formal academic training and more on intuitive moral reasoning tied to competition and authenticity. Future research should investigate whether students from other disciplines, or individuals with direct experience of enhancement technologies, show greater differentiation in their safety judgments.

Our findings also carry implications for regulation. Direct-to-consumer brain stimulation devices, for instance, are marketed as legal and non-invasive yet are received with moral ambivalence by students. This indicates that public acceptance cannot be assumed on the basis of legality alone. Regulators must therefore move beyond binary classifications of legal/illegal and safe/unsafe, toward pluralistic frameworks that account for fairness, authenticity, and identity concerns (Wexler, 2015). While much of the regulatory debate on direct-to-consumer brain stimulation has centred on the United States (Wexler, 2015), similar issues are beginning to surface in the UK and EU, where consumer neurotechnology is largely unregulated and falls between medical and consumer product frameworks. This regulatory uncertainty reflects a broader ethical tension between precautionary and proactionary approaches to emerging technologies (Bostrom & Sandberg, 2009): whether to restrict enhancement on the basis of potential risks and moral unease, or to permit innovation unless concrete harms are demonstrated. Our findings suggest that student attitudes resonate more strongly with precautionary intuitions, emphasising fairness, authenticity, and identity over legality or formal safety. Future policy debates will therefore need to balance these competing orientations, ensuring that regulation does not merely track legality or technical risk but also addresses the social and moral concerns that shape public legitimacy.

Several limitations should be considered when interpreting these findings. The study relied on self-report questionnaires, which may be influenced by social desirability and may not fully capture real-world behaviour or willingness to use enhancement technologies. The sample was drawn from a single UK university and consisted primarily of female psychology students, which restricts the generalisability of the results to broader or more diverse populations. The scenarios used to describe enhancement methods, while designed to be clear and balanced, cannot reflect the full complexity of lived decision-making. The cross-sectional design also prevents conclusions about how attitudes may shift over time or in response to direct experience with enhancement. Finally, the study examined three categories of enhancers and five moral domains, which provides an informative but incomplete picture of the broader ethical terrain.

Future research should address these limitations by recruiting more diverse and cross-cultural samples, including groups such as clinicians, policymakers, and working professionals who are directly involved in debates about enhancement. Longitudinal and experimental approaches could clarify how moral judgments change as technologies like brain stimulation become more familiar or as individuals acquire direct experience with them. Further work could also investigate how individual moral foundations, such as fairness, harm, and purity, predict differences in moral evaluation, and whether targeted interventions or public engagement can reshape perceptions. For the field of neuroethics, a particularly valuable direction will be to integrate empirical findings with normative analysis, examining not only how people evaluate enhancement but also whether and how those intuitions should guide ethical and regulatory frameworks. In this way, future work can help bridge descriptive accounts of moral psychology with prescriptive debates about the legitimate role of enhancement in education, healthcare, and society.

In summary, this study shows that attitudes toward cognitive enhancement are shaped not only by the type of method but also by how it is framed and situated within a social context. Brain stimulation, despite being legal and non-invasive, was judged more like pharmacological enhancement than natural methods, highlighting that moral evaluations are not reducible to risk or legality. Framing enhancement as preservation increased acceptability, while priming participants to consider competitive consequences reduced it, particularly for artificial methods. Together, these findings underscore the importance of integrating empirical insights into moral psychology with normative analysis in order to inform ethical debate and policy. Understanding how people evaluate enhancement in everyday academic life provides a crucial foundation for broader neuroethical discussions about fairness, authenticity, and the limits of self-improvement.

## Data Availability

Data will be available on request.

## Acknowledgements

We would like to thank all participants for their time and efforts.

## References

Bikson, M., Grossman, P., Thomas, C., Zannou, A. L., Jiang, J., Adnan, T., Mourdoukoutas, A. P., Kronberg, G., Truong, D., Boggio, P., Brunoni, A. R., Charvet, L., Fregni, F., Fritsch, B., Gillick, B., Hamilton, R. H., Hampstead, B. M., Jankord, R., Kirton, A., … Woods, A. J. (2016). Safety of transcranial Direct Current Stimulation: Evidence Based Update 2016. Brain Stimulation, 9(5), 641–661. 10.1016/j.brs.2016.06.004

Bostrom, N., & Sandberg, A. (2009). Cognitive enhancement: Methods, ethics, regulatory challenges. Science and Engineering Ethics, 15(3), 311–341. 10.1007/s11948-009-9142-5

Cabrera, L. Y., & Reiner, P. B. (2015). Understanding public (mis)understanding of tDCS for enhancement. Frontiers in Integrative Neuroscience, 9. 10.3389/fnint.2015.00030

Caviola, L., & Faber, N. S. (2015). Pills or Push-Ups? Effectiveness and Public Perception of Pharmacological and Non-Pharmacological Cognitive Enhancement. Frontiers in Psychology, 6, 1852. 10.3389/fpsyg.2015.01852

Cohen Kadosh, R., Levy, N., O’Shea, J., Shea, N., & Savulescu, J. (2012). The neuroethics of non-invasive brain stimulation. Current Biology: CB, 22(4), R108–R111. 10.1016/j.cub.2012.01.013

Conrad, E. C., Humphries, S., & Chatterjee, A. (2019). Attitudes Toward Cognitive Enhancement: The Role of Metaphor and Context. AJOB Neuroscience, 10(1), 35–47. 10.1080/21507740.2019.1595771

Costantino, A., Maiese, A., Lazzari, J., Casula, C., Turillazzi, E., Frati, P., & Fineschi, V. (2023). The Dark Side of Energy Drinks: A Comprehensive Review of Their Impact on the Human Body. Nutrients, 15(18), 3922. 10.3390/nu15183922

Dinh, C. T., Humphries, S., & Chatterjee, A. (2020). Public Opinion on Cognitive Enhancement Varies across Different Situations. AJOB Neuroscience, 11(4), 224–237. 10.1080/21507740.2020.1811797

Faul, F., Erdfelder, E., Lang, A.-G., & Buchner, A. (2007). G*Power 3: A flexible statistical power analysis program for the social, behavioral, and biomedical sciences. Behavior Research Methods, 39(2), 175–191. 10.3758/bf03193146

Fitz, N. S., Nadler, R., Manogaran, P., Chong, E. W. J., & Reiner, P. B. (2014). Public attitudes toward cognitive enhancement. Neuroethics, 7(2), 173–188. 10.1007/s12152-013-9190-z

Forlini, C., & Racine, E. (2009). Autonomy and Coercion in Academic “Cognitive Enhancement” Using Methylphenidate: Perspectives of Key Stakeholders. Neuroethics, 2(3), 163–177. 10.1007/s12152-009-9043-y

Franke, A. G., Christmann, M., Bonertz, C., Fellgiebel, A., Huss, M., & Lieb, K. (2011). Use of Coffee, Caffeinated Drinks and Caffeine Tablets for Cognitive Enhancement in Pupils and Students in Germany. Pharmacopsychiatry, 44, 331–338. 10.1055/s-0031-1286347

Fukuyama, F. (2002). Our posthuman future: Consequences of the biotechnology revolution. Farrar, Straus and Giroux.

Furrer, R. A., Merner, A. R., Stevens, I., Zuk, P., Williamson, T., Shen, F. X., & Lázaro-Muñoz, G. (2025). Public perceptions of neurotechnologies used to target mood, memory, and motor symptoms. Device, 100804. 10.1016/j.device.2025.100804

Grinschgl, S., Ninaus, M., Wood, G., & Neubauer, A. C. (2025). To enhance or not to enhance: A debate about cognitive enhancement from a psychological and neuroscientific perspective. Physics of Life Reviews, 54, 58–77. 10.1016/j.plrev.2025.05.002

Maier, L. J., Liakoni, E., Schildmann, J., Schaub, M. P., & Liechti, M. E. (2015). Swiss University Students’ Attitudes toward Pharmacological Cognitive Enhancement. PloS One, 10(12), e0144402. 10.1371/journal.pone.0144402

Maslen, H., Earp, B. D., Kadosh, R. C., & Savulescu, J. (2014). Brain stimulation for treatment and enhancement in children: An ethical analysis. Frontiers in Human Neuroscience, 8. 10.3389/fnhum.2014.00953

Mayor, E., Daehne, M., & Bianchi, R. (2019). How perceived substance characteristics affect ethical judgement towards cognitive enhancement. PLOS ONE, 14(3), e0213619. 10.1371/journal.pone.0213619

McDermott, H., Lane, H., & Alonso, M. (2021). Working smart: The use of ‘cognitive enhancers’ by UK university students. Journal of Further and Higher Education, 45(2), 270–283. 10.1080/0309877X.2020.1753179

Medaglia, J. D., Yaden, D. B., Helion, C., & Haslam, M. (2019). Moral attitudes and willingness to enhance and repair cognition with brain stimulation. Brain Stimulation, 12(1), 44–53. 10.1016/j.brs.2018.09.014

Mihailov, E., Rodríguez López, B., Cova, F., & Hannikainen, I. R. (2021). How pills undermine skills: Moralization of cognitive enhancement and causal selection. Consciousness and Cognition, 91, 103120. 10.1016/j.concog.2021.103120

Parens, E. (2005). Authenticity and Ambivalence: Toward Understanding the Enhancement Debate. The Hastings Center Report, 35(3), 34–41. 10.2307/3528804

Patton, H. (2023). Development of the Attitudes Towards Pharmacological Cognitive Enhancement Scale. Trent University.

Schelle, K. J., Faulmüller, N., Caviola, L., & Hewstone, M. (2014). Attitudes toward pharmacological cognitive enhancement—A review. Frontiers in Systems Neuroscience, 8, 53. 10.3389/fnsys.2014.00053

Schermer, M. (2008). On the argument that enhancement is ‘cheating’. Journal of Medical Ethics, 34(2), 85–88.

Scheske, C., & Schnall, S. (2012). The Ethics of “Smart Drugs”: Moral Judgments About Healthy People’s Use of Cognitive-Enhancing Drugs. Basic and Applied Social Psychology, 34(6), 508–515. 10.1080/01973533.2012.711692

Veit, W., Earp, B. D., Faber, N., Bostrom, N., Caouette, J., Mannino, A., Caviola, L., Sandberg, A., & Savulescu, J. (2020). Recognizing the Diversity of Cognitive Enhancements. AJOB Neuroscience, 11(4), 250–253. 10.1080/21507740.2020.1830878

Wexler, A. (2015). A pragmatic analysis of the regulation of consumer transcranial direct current stimulation (TDCS) devices in the United States. Journal of Law and the Biosciences, 2(3), 669–696. 10.1093/jlb/lsv039

