## Supplemental Table 1 for "Legal but immoral: attitudes toward non-invasive brain stimulation for cognitive enhancement"

**Supplementary 1.**

**Framing scenarios**

*Enhance*

Imagine a university student preparing for final exams and wanting to enhance their focus and memory. They decide to start drinking coffee daily. After several days, the student notices they can study for longer periods without losing concentration and solve problems more effectively. As a result, they performed 15% better on practice tests and felt more confident heading into exams.

Imagine a university student preparing for final exams and wanting to enhance their focus and memory. They purchase a legal, non-invasive brain stimulation headset that boosts attention and cognitive function. This works by delivering mild electrical currents to the prefrontal cortex, which stimulates neurons. After several days, the student notices they can study for longer periods without losing concentration and solve problems more effectively. As a result, they performed 15% better on practice tests and felt more confident heading into exams. The student had no reported adverse side effects.

Imagine a university student preparing for final exams and wanting to enhance their focus and memory. They begin taking medication (similar to Adderall or Ritalin that was not prescribed to them) that boosts attention and cognitive function by increasing neurotransmitter activity. After several days, the student notices they can study for longer periods without losing concentration and solve problems more effectively. As a result, they performed 15% better on practice tests and felt more confident heading into exams. The student had no reported adverse side effects.

*Preserve*

Imagine a university student who is burnt out and mentally fatigued. They are preparing for final exams and want to preserve their focus and memory. They decide to start drinking coffee daily. After several days, the student notices they can study for longer periods without losing concentration and solve problems more effectively. As a result, they performed 15% better on practice tests and felt more confident heading into exams.

Imagine a university student who is burnt out and mentally fatigued. They are preparing for final exams and want to preserve their focus and memory. They purchase a legal, non-invasive brain stimulation headset that boosts attention and cognitive function. This works by delivering mild electrical currents to the prefrontal cortex, which stimulates neurons. After several days, the student notices they can study for longer periods without losing concentration and solve problems more effectively. As a result, they performed 15% better on practice tests and felt more confident heading into exams. The student had no reported adverse side effects.

Imagine a university student who is burnt out and mentally fatigued. They are preparing for final exams and want to preserve their focus and memory. They begin taking a medication (similar to Adderall or Ritalin that was not prescribed to them) that boosts attention and cognitive function by increasing neurotransmitter activity. After several days, the student notices they can study for longer periods without losing concentration and solve problems more effectively. As a result, they performed 15% better on practice tests and felt more confident heading into exams. The student had no reported adverse side effects.

**Priming**

The Bell Curve produces a normal distribution of grades at university. A small percentage of students are deemed able to achieve First-Class Honours (1st), whereas most other students are assigned lower grades, such as Upper Second-Class Honours (2:1) and Lower Second-Class Honours (2:2). This means that the performance of other students has a direct impact on your position on the Bell Curve, therefore impacting your final grade.

Grades at university are distributed in a manner where any percentage of students are able to achieve First-Class Honours (1st), regardless of the percentage of other students who achieve average grades of Upper Second-Class Honours (2:1) and Lower Second-Class Honours (2:2). This means that the performance of other students has no direct impact on your final grade.
